# Block HSIC Lasso: model-free biomarker detection for ultra-high dimensional data

**DOI:** 10.1101/532192

**Authors:** Héctor Climente-González, Chloé-Agathe Azencott, Samuel Kaski, Makoto Yamada

**Affiliations:** Institut Curie, PSL Research University, F-75005 Paris, France; INSERM, U900, F-75005 Paris, France; MINES ParisTech, PSL Research University, CBIO-Centre for Computational Biology, F-75006 Paris, France; Aalto University, Helsinki, Finland; Kyoto University, Kyoto, 606-8501, Japan; RIKEN AIP, Chuo-ku, Tokyo 103-0027, Japan

## Abstract

**Motivation:** Finding nonlinear relationships between biomolecules and a biological outcome is computationally expensive and statistically challenging. Existing methods have important drawbacks, including among others lack of parsimony, non-convexity, and computational overhead. Here we propose block HSIC Lasso, a nonlinear feature selector that does not present the previous drawbacks.

**Results:** We compare block HSIC Lasso to other state-of-the-art feature selection techniques in both synthetic and real data, including experiments over three common types of genomic data: gene-expression microarrays, single-cell RNA sequencing, and genome-wide association studies. In all cases, we observe that features selected by block HSIC Lasso retain more information about the underlying biology than those selected by other techniques. As a proof of concept, we applied block HSIC Lasso to a single-cell RNA sequencing experiment on mouse hippocampus. We discovered that many genes linked in the past to brain development and function are involved in the biological differences between the types of neurons.

**Availability:** Block HSIC Lasso is implemented in the Python 2/3 package pyHSICLasso, available on PyPI. Source code is available on GitHub (https://github.com/riken-aip/pyHSICLasso).

**Supplementary information:** Supplementary data are available at *Bioinformatics* online.

## 1 Introduction

Biomarker discovery, the goal of many bioinformatics experiments, aims at identifying a few key biomolecules that explain most of an observed phenotype. Without a strong prior hypothesis, these molecular markers have to be identified from data generated by high-throughput technologies. Unfortunately, finding relevant molecules is a combinatorial problem: for *d* features, 2^*d*^ binary choices must be considered. As the number of features vastly exceeds the number of samples, biomarker discovery is a high-dimensional problem. The statistical challenges posed by such high-dimensional spaces have been thoroughly reviewed elsewhere (Clarke *et al*., 2008; Johnstone and Titterington, 2009). In general, due to the *curse of dimensionality*, fitting models in many dimensions and on a small number of samples is extremely hard. Moreover, since biology is complex, a simple statistical model such as a linear regression might not be able to find important biomarkers. Those that are found in such experiments are often hard to reproduce, suggesting overfitting. Exploring the solution space and finding true biomarkers is not only statistically challenging, but also computationally expensive.

In machine learning terms, biomarker discovery can be formulated as a problem of feature selection: identifying the best subset of features to separate between categories, or to predict a continuous response. In the past decades, many feature selection algorithms that deal with high-dimensional datasets have been proposed. Due to the difficulties posed by high-dimensionality, linear methods tend to be the feature selector of choice in bioinformatics. A widely used linear feature selector is the Least Absolute Shrinkage and Selection Operator, or Lasso (Tibshirani, 1996). Lasso fits a linear model between the input features and phenotype by minimizing the sum of the least square loss and an *ℓ*_1_ penalty term. The balance between the least square loss and the penalty ensures that the model explains the linear combination of features, while keeping the number of features in the model small. However, in many instances biological phenomena do not behave linearly. In such cases there is no guarantee that Lasso can capture those nonlinear relationships or an appropriate effect size to represent them.

In the past decade, several nonlinear feature selection algorithms for high-dimensional datasets have been proposed. One of the most widely used, called Sparse Additive Model, or SpAM (Ravikumar *et al*., 2009), models the outcome as a sparse linear combination of nonlinear functions based on kernels. However, since SpAM assumes an additive model over the selected features, it cannot select important features if the phenotype cannot be represented by the additive functions of input features – for example, if there exist a multiplicative relationship between features (Yamada *et al*., 2014).

Another family of nonlinear feature selectors are association-based: they compute the statistical association score between each input feature and the outcome, and rank features accordingly. Since these approaches do not assume any model about the output, they can detect important features as long as an association exists. When using a nonlinear association measure, such as the mutual information (Cover and Thomas, 2006) or the Hilbert-Schmidt Independence Criterion (HSIC) (Gretton *et al*., 2005), they select the features with the strongest dependence with the phenotype. However, association-based methods do not account for the redundancy between the features, which is frequent in biological data sets, since they do not model relationships between features. Hence, many redundant features are typically selected, hindering interpretability. This is important in applications like drug target discovery, where only a small number of targets can be validated, and it is crucial to discriminate the most important target out of many other top-ranked targets.

To deal with the problem of redundant features, Peng *et al*. (2005) proposed the minimum redundancy maximum relevance (mRMR) algorithm. mRMR can select a set of non-redundant features that have high association to the phenotype, while penalizing the selection of mutually dependent features. Ding and Peng (2005) used mRMR to extract biomarkers from microarray data, finding that the selected genes captured better the variability in the phenotypes than those identified by state-of-the-art approaches. However, mRMR has three main drawbacks: the optimization problem is discrete; it must be solved by a greedy approach; and the mutual information estimation is difficult (Walters-Williams and Li, 2009). Moreover, it is unknown whether the objective function of mRMR has good theoretical properties such as submodularity (Fujishige, 2005), which would guarantee the optimality of the solution.

Recently, Yamada *et al*. (2014) proposed a kernel-based minimum redundancy maximum relevance algorithm called HSIC Lasso. Instead of mutual information, HSIC Lasso employs the Hilbert-Schmidt Independence Criterion (HSIC) (Gretton *et al*., 2005) to measure dependency between variables. In addition, it uses an *ℓ*_1_ penalty term to select a small number of features. This results in a convex optimization problem, for which one can therefore find a globally optimal solution. In practice, HSIC Lasso has been found to outperform mRMR in several experimental settings (Yamada *et al*., 2014). However, HSIC Lasso is memory-intensive: its memory complexity is *O*(*dn*^2^), where *d* is the number of features and *n* is the number of samples. Hence, HSIC Lasso cannot be applied to datasets with thousands of samples, nowadays widespread in biology. A MapReduce version of HSIC Lasso has been proposed to address this drawback, and it is able to select features in ultra-high dimensional settings (10^6^ features, 10^4^ samples) in a matter of hours (Yamada *et al*., 2018). However, it requires a large number of computing nodes, inaccessible to common laboratories. And, since it relies on the Nyström approximation of Gram matrices (Schölkopf and Smola, 2002), the final optimization problem is no longer convex, and hence finding a globally optimal solution cannot be easily guaranteed.

In this paper, we propose block HSIC Lasso: a simple yet effective nonlinear feature selection algorithm based on HSIC Lasso. The key idea is to use the recently proposed block HSIC estimator (Zhang *et al*., 2018) to estimate the HSIC terms. By splitting the data in blocks of size *B* ≪ *n*, the memory complexity of HSIC Lasso goes from *O*(*dn*^2^) down to *O*(*dnB*). Moreover the optimization problem of the block HSIC Lasso remains convex. Through its application to synthetic data and biological datasets, we show that block HSIC Lasso can be applied to a variety of settings and compares favorably with the vanilla HSIC Lasso algorithm and other feature selection approaches, linear and nonlinear, as it selects features more informative of the biological outcome. Further considerations on the state-of-the-art and the relevance of block HSIC Lasso can be found in Supplementary File 1.

## 2 Materials and methods

### 2.1 Problem formulation

Assume a data set with n samples described by *d* real-valued features, each corresponding to a biomolecule (for example, the expression of one transcript, or the number of major alleles observed at a given SNP), and a label, continuous or binary, describing the outcome of interest (for example, the abundance of a target protein, or disease status). We denote the *i*-th sample by 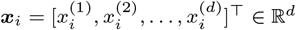, where ⊤ denotes transpose; and its label by 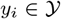, where 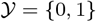 for a binary outcome, corresponding to a classification problem, and 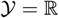 for a continuous outcome, corresponding to a regression problem. In addition, we denote by 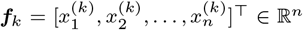 the *k*-th feature in the data.

The goal of supervised feature selection is to find *m* features (*m* ≪ *d*) that are the most relevant for predicting the output *y* for a sample ***x***.

### 2.2 HSIC Lasso

Measuring the dependence between two random variables *X* and *Y* can be achieved by the Hilbert-Schimdt Independence Criterion, or HSIC (Gretton *et al*., 2005):

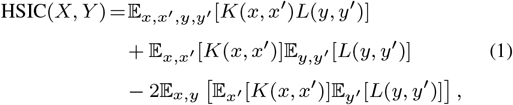

where *K*: ℝ^*d*^ × ℝ^d^ → ℝ and 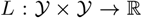 are positive definite kernels, and 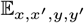 denotes the expectation over independent pairs (*x, y*) and (*x*′, *y*′) drawn from *p*(*x, y*). HSIC(*X, Y*) is equal to 0 if *X* and *Y* are independent, and is non-negative otherwise.

In practice, for a given Gram matrix ***K**_k_* ∈ ℝ^*n* × *n*^, computed from the *k*-th feature, and a given output Gram matrix **L** ∈ ℝ^*n*×*n*^, the normalized variant of HSIC is computed using its *V*-statistic estimator as (Yamada *et al*., 2018)

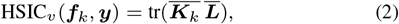

where for a Gram matrix ***K*** ∈ ℝ^*n*×*n*^, 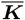 is defined as 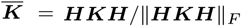 with ***H*** ∈ ℝ^*n*×*n*^ a centering matrix defined by 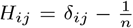. Here *δ_ij_* is equal to 1 if *i* = *j* and 0 otherwise, and tr denotes the trace. Note that we employ the normalized variant of the original empirical HSIC.

The largest the value of HSIC_*v*_ (***f**_k_, **y***), and the more dependent the *k*-th feature and the outcome are. Song *et al*. (2012) therefore proposed to perform feature selection by ranking the features by descending value of HSIC_*v*_(***f**_k_, **y***).

With HSIC Lasso, Yamada *et al*. (2014) extend the work of Song *et al*. (2012) so as to avoid selecting multiple redundant features. For this purpose, they introduce a vector ***α*** = [*α*_1_,…, *α_d_*]^⊤^ of feature weights and solve the following optimization problem:

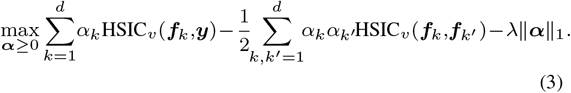

The first term enforces selected features that are highly dependent on the phenotype; the second term penalizes selecting mutually dependent features; and the third term enforces selecting a small number of features. The selected features are those that have a non-zero coefficient *α_k_*. Here *λ* > 0 is a regularization parameter that controls the sparsity of the solution: the larger *λ*, the fewer features have a non-zero coefficient.

The HSIC Lasso optimization problem can be rewritten as

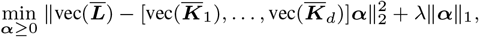

where vec: ℝ^*n*×*n*^ → ℝ^*n*^2^^, ***K*** ↦ [*K*_11_,…, *K*_1*n*_, *K*_21_,…, *K_nn_*] is the vectorization operator. Using this formulation, we can solve the problem using an off-the-shelf non-negative Lasso solver.

HSIC Lasso performs well for high-dimensional data. However, it requires a large memory space (*O*(*dn*^2^)), since it stores *d* Gram matrices. To handle this issue, two approximation methods have been proposed. The first approach uses a memory lookup to dramatically reduce the memory space (Yamada *et al*., 2014). However, since this method needs to perform a large number of memory lookups, it is computationally expensive. Another approach (Yamada *et al*., 2018) is to rewrite the problem using the Nyström approximation (Schölkopf and Smola, 2002) and solve the problem using a cluster. However using the Nyström approximation makes the problem non-convex.

### 2.3 Block HSIC Lasso

In this paper, we propose an alternative HSIC Lasso method for large-scale problems, the *block HSIC Lasso*, which is convex and can be efficiently solved on a reasonably-sized server.

Block HSIC Lasso employs the block HSIC estimator (Zhang *et al*., 2018) instead of the V-statistics estimator of Equation (2). More specifically, to compute the block HSIC, we first partition the training dataset into 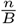 partitions 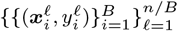, where *B* is the number of samples in each block. Note that the block size *B* is set to a relatively small number such as 10 or 20 (*B* ≪ *n*). Then, the block HSIC estimator can be written as

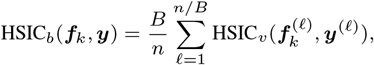

where 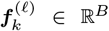 represents the *k*-th feature vector of the *ℓ*-th partition. Note that the computation of 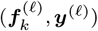 requires *O*(*B*^2^) memory space. Therefore, the required memory for the block HSIC estimator is *O*(*nB*^2^), where *nB* ≪ *n*^2^.

If we denote by 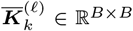 the restriction of 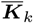 to the *ℓ*-th partition, and by 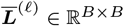 the restriction of ***L*** to the *ℓ*-th partition, then

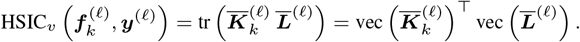

Block HSIC Lasso is obtained by replacing the HSIC estimator HSIC_*v*_ with the block HSIC estimator HSIC_*b*_ in Equation (3):

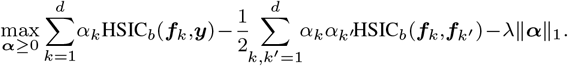

Using the vectorization operator, the block estimator is written as

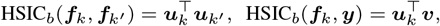

where

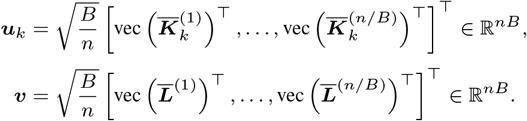

Hence, block HSIC Lasso can also be written as

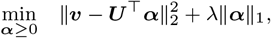

where ***U*** = [***u***_1_,…,***u***_*d*_] ∈ ℝ^*nB*×*d*^.

Since the objective function of block HSIC Lasso is convex, we can obtain a globally optimal solution. As with HSIC Lasso, we can solve block HSIC Lasso using an off-the-shelf Lasso solver. Here, we use the non-negative least angle regression-LASSO, or LARS-LASSO (Efron *et al*., 2004), to solve the problem in a greedy manner. Rather than setting the hyperparameter *λ*, for example by cross-validation, which would be computationally intensive, this allows us to use a predefined number of features to select.

The required memory space for block HSIC Lasso is *O*(*dnB*), which compares favorably to vanilla HSIC Lasso’s *O*(*dn*^2^); as the block size *B* ≪ *n*, the memory space is dramatically reduced. However, the computational cost of the proposed method is still large when both *d* and *n* are large. Thus, we implemented the proposed algorithm using multiprocessing by parallelizing the computation of 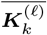. Thanks to the combination of block HSIC Lasso and the multiprocessing implementation, we can efficiently find solutions on large datasets with a reasonably-sized server.

### 2.4 Improving selection stability using bagging

Since we need to compute block HSIC of the paired data 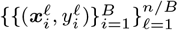 with a fixed partition, the performance can be highly affected by the partition. Thus, we propose to use a bagging version of the block HSIC estimator. Given *M* random permutations of the *n* samples, we define *bagging block HSIC* as

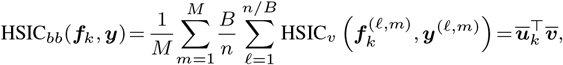

where 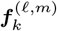 is the *k*-th feature vector restricted to the *ℓ*-th block as defined by the *m*-th permutation,

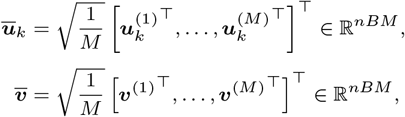

and 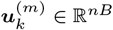 and 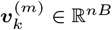 are the vectors of the *m*-th block HSIC Lasso, respectively.

Hence, bagging block HSIC Lasso can be written as

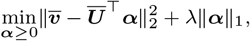

where 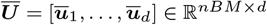.

We consider the bagging part to be an integral part of the block HSIC Lasso algorithm. That is why, in this text, every time we mention “block HSIC Lasso”, we refer to bagging block HSIC Lasso.

Note that the memory space *O*(*dnBM*) required for *B* = 60 and *M* = 1 is equivalent to *B* = 30 and *M* = 2. Empirically, we found that they were providing equivalent feature selection accuracy (Section 4.4).

### 2.5 Adjusting for covariates

Data analysis tasks in bioinformatics can often be confounded by technical (e.g. batch) or biological variables (e.g. age), which might mask the relevant variables. To adjust for their effect, we consider the following variant of the block HSIC Lasso:

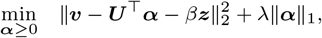

where *β* ≥ 0 is a tuning parameter and

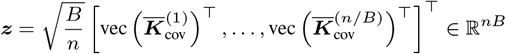

contains the covariate information. ***K***_cov_ is the Gram matrix computed from the covariate input matrix ***X***_cov_. Since for most purposes in bioinformatics we want to remove all information from the covariates, we set *β* to

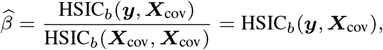

which is the solution of 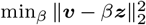. Here, we used the property HSIC_*b*_(***X***_cov_, ***X***_cov_) = 1.

## 3 Experimental setup

### 3.1 Feature selection methods

#### HSIC Lasso and block HSIC Lasso

We used HSIC Lasso and block HSIC Lasso implemented in the Python 2/3 package *pyHSICLasso*. In block HSIC Lasso, *M* was set to 3 in all experimental settings; the block size *B* was set on an experiment-dependent fashion. In all the experiments, when we wanted to select *k* features, HSIC Lasso versions were required to first retrieve 50 features, and then the top *k* features were selected as the solution.

In this paper, we use the following kernels:

- The RBF Gaussian kernel for pairs of continuous variables, of continuous outcomes, or one of each, and for pairs of a continuous variable and categorical outcome:

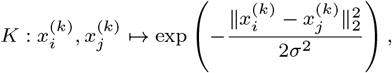

where *σ*^2^ > 0 is the bandwidth of the kernel;
- The normalized Delta kernel for categorical variables (or outcomes):

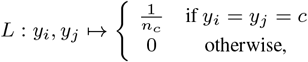

where *n_c_* is the number of samples in class *c*.

#### mRMR

Minimum Redundancy Maximum Relevance (mRMR) selects features that are highly associated with the outcome and are non-redundant (Peng *et al*., 2005). To that end, it uses mutual information between different variables and between the outcome and the variables.

We used a C++ implementation of mRMR (Peng, 2005). The maximum number of samples and the maximum number of features were set to the actual number of samples and features in the data. In regression problems, discretization was set to binarization.

#### LARS

Least angle regression (LARS) is a forward stagewise feature selector (Efron *et al*., 2004). It is an efficient way of solving the same problem as Lasso. We used the SPAMS implementation of LARS (Mairal *et al*., 2010), with the default parameters. Note that this is not the implementation of LARS that we use in (block) HSIC Lasso, which is the nonnegative LARS solver implemented in *pyHSICLasso*.

### 3.2 Evaluation of the selected features

#### Selection accuracy on simulated data

We simulated highdimensional data where only a few variables were truly related to the outcome. We used these datasets to evaluate the ability of the tested algorithms to find the true causal variables, instead of others, likely spuriously correlated to the outcome. To that end, we requested each algorithm to retrieve the known number of causal features. Then, we studied how many of them were actually causal.

#### Classification with a random forest

In classification data sets, we evaluated the amount of information retained in the features selected by a given method by evaluating the performance of a random forest classifier based only on those features. We used random forests because their ability to handle nonlinearities. We split the data between a training and a test set, and selected features on the training set only. We estimated the best parameters by cross-validation on the training set: the number of trees (200, 500), the maximum depth of the threes (4, 6, 8), the number of features to consider 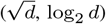, and the criterion to measure the quality of the chosen features (Gini impurity, information gain). Then, we trained a model with those parameters on the training set and made predictions on a separate testing set to estimate prediction accuracy.

### 3.3 Datasets

We evaluated the performance of the different algorithms on synthetic data and four types of real-world high dimensional datasets (Table 1). In our experiments on real-world datasets, we restricted ourselves to classification problems. All discussed methods can however handle regression problems (continuous-valued outcomes) as well, as we show on synthetic data.

**Table 1.**
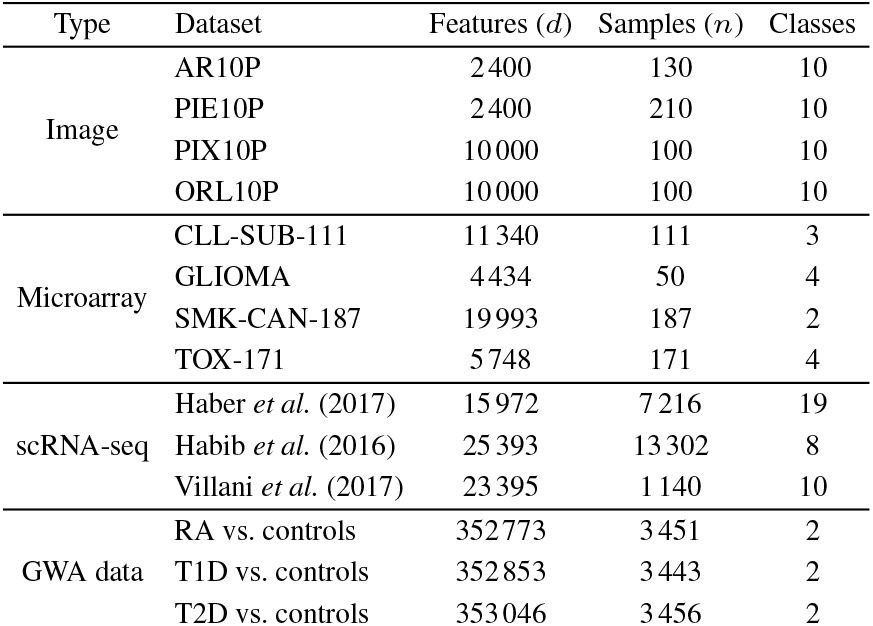
Summary description of benchmark datasets.

#### Synthetic data

We simulated random matrices of features 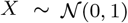. A number of variables were selected as related to the phenotype, and functions that are nonlinear in the data range were selected (cosine, sine and square) and combined additively to create the outcome vector ***y***.

#### Images

Facial recognition is a classification problem classically used to evaluate nonlinear feature selection methods, as only a few of all features are expected to be relevant for the outcome, in a nonlinear fashion. We used four face image datasets from the Arizona State University feature selection repository (Li *et al*., 2018)): pixraw10P, warpAR10P, orlraws10P, and warpPIE10P.

#### Gene expression microarrays

We analyzed four gene expression microarray datasets from Arizona State University feature selection repository (Li *et al*., 2018). The phenotypes were subtypes of B-cell chronic lymphocytic leukemia (CLL-SUB-111), hepatocyte phenotypes under different diets (TOX-171), glioma (GLIOMA) and smoking-driven carcinogenesis (SMK-CAN-187).

#### Single-cell RNA-seq

Single-cell RNA-seq (scRNA-seq) measures gene expression at cell resolution, allowing to characterize the diversity in a tissue. We performed feature selection on the three most popular datasets in the Broad Institute’s Single Cell Portal, related to mouse small intestinal epithelium (Haber *et al*., 2017), mouse hippocampus (Habib *et al*., 2016), and human blood cells (Villani *et al*., 2017). Missing gene expressions were imputed with MAGIC (van Dijk *et al*., 2018)).

#### GWA datasets

We studied the WTCCC1 datasets (Burton *et al*., 2007) for rheumatoid arthritis (RA), type 1 diabetes (T1D) and type 2 diabetes (T2D) (2000 samples each), using the 1958BC cohort as control (1504 samples). Affymetrix 500K was used for genotyping. We removed the samples and the SNPs that did not pass WTCCC’s quality controls, as well as SNPs in sex chromosomes and those that were not genotyped in both cases and controls. Missing genotypes were imputed with CHIAMO. Lastly, individuals with >10% genotype missing rate, and SNPs with >10% genotype missing rate, MAF < %5 or not in HWE (p-value < 0.001) were removed. The remaining missing genotypes were replaced by the major allele in homozygosis.

#### Preprocessing

Images, microarrays and scRNA-seq data were normalized feature-wise by subtracting the mean and dividing by the standard deviation. GWAS data did not undergo any normalization.

### 3.4 Computational resources

We ran the experiments on synthetic data, images, microarrays and scRNA-seq on CentOS 7 machines with Intel Xeon 2.6GHz and 50GB RAM memory. For the GWA datasets experiments, we used a CentOS 7 server with 96 core Intel Xeon 2.2GHz and 1TB RAM memory.

### 3.5 Software availability and reproducibility

Block HSIC Lasso was implemented in the Python 2/3 package *pyHSICLasso*. The source code is available on GitHub (https://github.com/riken-aip/pyHSICLasso), and the package can be installed from PyPI (https://pypi.org/project/pyHSICLasso). All analyses in this paper and the scripts needed to reproduce them are also available on GitHub (https://github.com/hclimente/nori).

## 4 Results

### 4.1 Block HSIC Lasso performance is comparable to state of the art

At first, we worked on synthetic, nonlinear data (section 3.2). We generated synthetic data with combinations of the following experimental parameters: *n* = {100, 1 000, 10 000} samples; *d* = {100, 2 500, 5 000, 10 000} features; and 5, 10 and 20 causal features, that is, features truly related to the outcome. We evaluated the performance of different feature selectors at retrieving the causal features. These conditions range from an ideal setting, where the number of features is smaller than the number of samples, to an ultra-high dimensional scenario, where spurious dependencies among variables, and between those and the outcome are bound to occur.

Each of the methods was required to select as many features as the number of true causal features. In Figure 1 we show the proportion of the causal features retrieved by each method. The different versions of HSIC Lasso outperform the other approaches in virtually all settings. Block HSIC Lasso with decreasing block sizes results in worse performances. As expected, vanilla HSIC Lasso outperforms the block versions in accuracy, but increases memory use. Crucially, block HSIC Lasso on a larger number of samples performs better than vanilla HSIC Lasso on fewer samples. Hence, when the number of samples is in the thousands, it is better to apply block HSIC Lasso on the whole dataset, than to apply vanilla HSIC Lasso on a subsample.

**Fig. 1.**
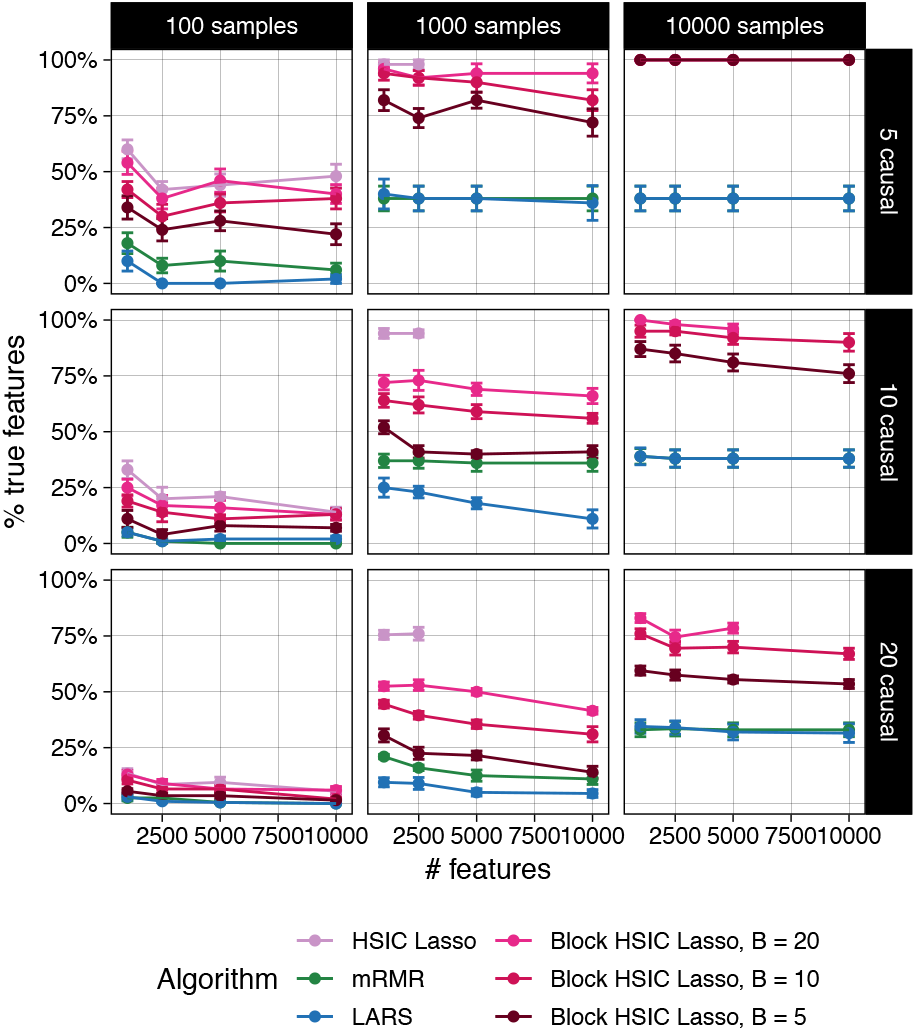
Percentage of true causal features extracted by different feature selectors. Each datapoint represents the mean over 10 replicates, and the error bars represent the standard error of the mean. Lines are discontinued when the algorithm required more memory than the provided (50 GB). Note that in some conditions mRMR’s line cannot be seen due to the overlap with LARS.

We wanted to test these conclusions using a nonlinear, real-world dataset. We selected four image-based face recognition tasks (Section 3.3). In this case, we selected different numbers of features (10, 20, 30, 40 and 50). Then, we trained random forest classifiers on these subsets of the features, and compared the accuracy of the different classifiers on a test set (Figure S1). Block HSIC Lasso displayed a performance comparable to vanilla HSIC Lasso, and comparable or superior to the other methods. This is remarkable, since it shows that, in many practical cases, block HSIC Lasso does not need more samples to achieve vanilla HSIC Lasso performance.

### 4.2 Adjusting by covariates improves feature selection

To evaluate the impact of covariate adjustment, we worked on a synthetic dataset (Section 3.2) with the following experimental parameters: *n* = 1 000; *d* = {100, 2 500, 5 000, 10 000} features; 7 causal features. Two covariates were generated by taking two causal features and adding Gaussian noise (mean = 0; standard deviation = 0.5). In the experiment shown in Figure S2, we tested the ability of (block) HSIC Lasso to retrieve exclusively the remaining 5 causal features adjusting for the covariates. We observe that block HSIC Lasso is able to find more relevant features when it adjusts for known covariates.

### 4.3 Block HSIC Lasso is computationally efficient

In our experiments on synthetic data, vanilla HSIC Lasso runs into memory issues already with 1 000 samples (Figure 1). This experiment shows how block HSIC Lasso keeps the good properties of HSIC Lasso, while extending it to more experimental settings. Block HSIC Lasso with *B* = 20 reaches the memory limit only at 10 000 samples, which is already sufficient for most common bioinformatics applications. If larger datasets need to be handled, that can be done by using smaller block sizes or a larger computer cluster.

We next quantified the computational efficiency improvement the block HSIC estimator brings. We compared the runtime and the peak memory usage in the highest-dimensional setting where all methods could run (*n* = 1000, *d* = 2500, 20 causal features) (Figure 2). We observe how, as expected, block HSIC Lasso requires an order of magnitude less memory than vanilla HSIC Lasso. Block versions also run notoriously faster, thanks to the lower number of operations and the parallelization. mRMR is ten times faster than block HSIC Lasso, at the expense of a clearly lower accuracy. However, a fraction of this gap is likely due to mRMR having been implemented in C++, while HSIC Lasso is written in Python. In this regard, there is potential for other faster implementations of (block) HSIC Lasso.

**Fig. 2.**
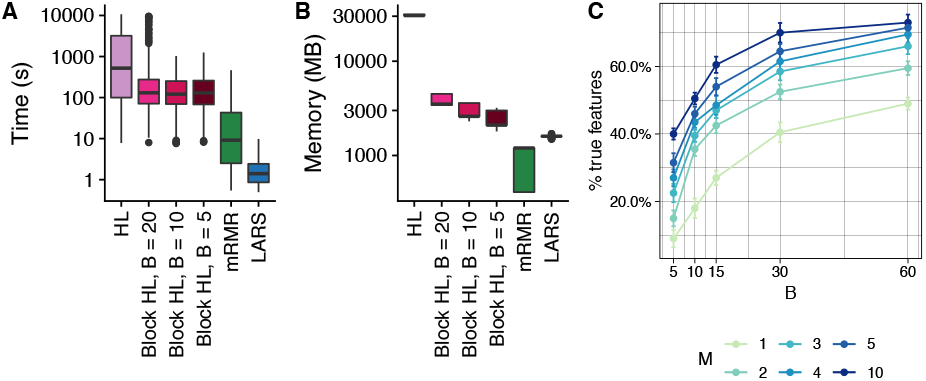
Computational resources used by the different methods. A. Time elapsed in a multiprocess setting. B. Memory usage in a single-core setting. C. Number of correct features retrieved on synthetic data (*n* = 1000, *d* = 2500, 20 causal features) by block HSIC Lasso at different block sizes *B* and number of permutations *M*.

### 4.4 Block HSIC Lasso improves with more permutations

We were interested in the trade-off between the block size and the number of permutations, which affect both the computation time and accuracy of the result. We tested the performance of block HSIC Lasso with *B* = {5, 10, 15, 30, 60} and *M* = {1, 2, 3, 5} in datasets of *n* = 1000, *d* = 2500 and 20 causal features. As expected, causal feature recovery increases with *M* and *B* (Figure 2C), as the HSIC estimator approaches its true value.

The memory usage *O*(*dnBM*) of several of the conditions was the same e.g. *B* =10, *M* = 3 and *B* = 30, *M* =1. Such conditions are indistinct from the points of view of both accuracy, and memory requirements. In practice, we found no major differences in runtime between different combinations of *B* and *M*. Hence, a reasonable strategy is to fix *B* to a given size, and tune the *M* to the available memory/desired amount of information. This strategy, however, should be adapted to fit properties of the data. More specifically, GWAS data is notably sparse, and as result a small block size would result in many blocks consisting entirely of zeros, which would hence be uninformative. In such cases it might be interesting to prioritize larger block sizes, and fewer permutations.

### 4.5 Block HSIC Lasso finds more relevant features

We tested the dimensionality reduction potential of different feature selectors. We selected a variable number of features from different multiclass biological datasets, then used a random forest classifier to retrieve the original classes (Section 3.2). The underlying assumption is that only selected features which are biologically relevant will be useful to classify unseen data. To that end, we evaluated the classification ability of the biomarkers selected in four gene expression microarrays (Figure 3) and three scRNA-seq experiments (Figure S3). Unsurprisingly, we observe that nonlinear feature selectors perform notably better than linear selectors. Of the nonlinear methods, in virtually all cases block HSIC Lasso showed similar or superior performance to mRMR. Interestingly, as little as 20 selected genes retain enough information to achieve a plateau accuracy in most experiments.

**Fig. 3.**
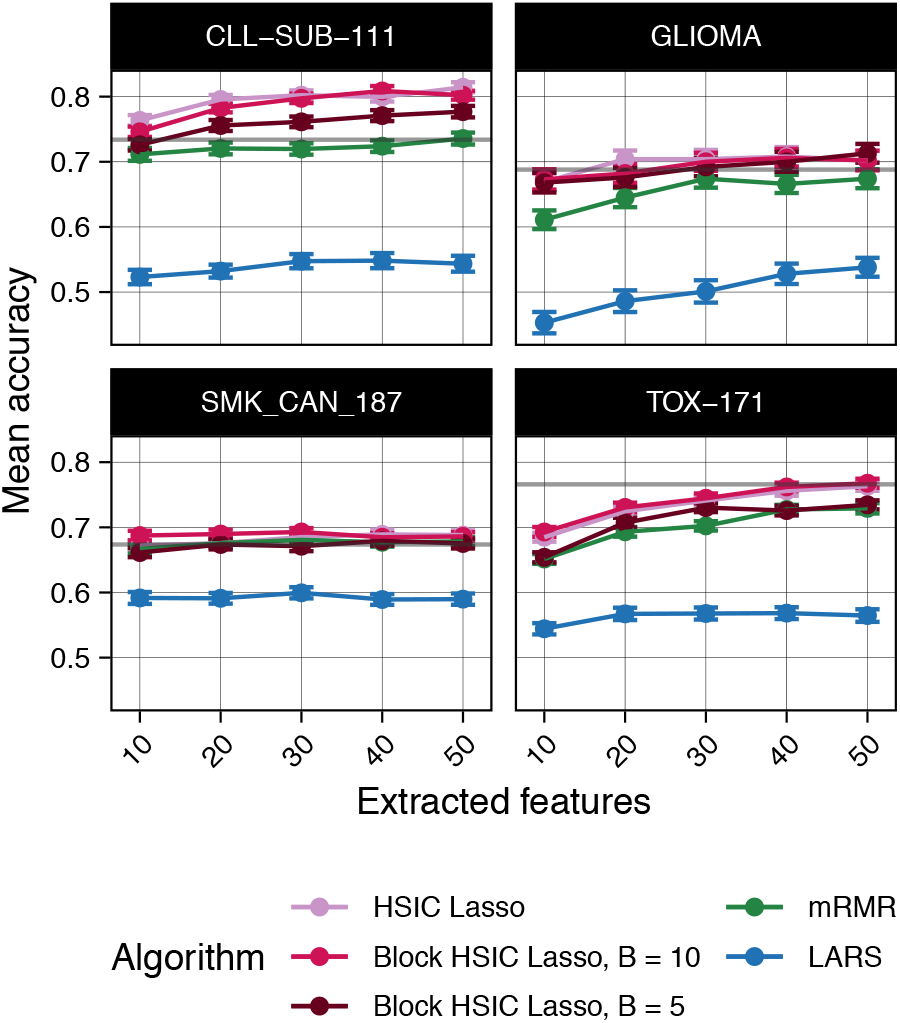
Random forest classification accuracy of microarray gene expression samples after feature extraction by the different methods. The gray line represents the mean accuracy of 10 classifiers trained on all the dataset.

Surveying 10^5^ – 10^6^ SNPs in 10^3^ – 10^4^ patients, genome-wide association (GWA) datasets are among the most high-dimensional in biology, an unbalance which worsens the statistical and computational challenges. We performed the same evaluation on three WTCCC1 phenotypes (Section 3.3). As a baseline, we also computed the accuracy of a classifier trained on all the SNPs (Table S1). We observe that a feature selection prior step is not always favourable: LARS worsens the classification accuracy by 5-10%. On top of that, LARS could not select any SNP in 2 out of the 15 experimental settings. On the other hand, nonlinear methods improve the classification accuracy by 10%, with mRMR and block HSIC Lasso achieving similar accuracies. In fact, those two selected the same 14 out of 30 SNPs when we selected 10 SNPs in each the three datasets with each method (Figure 7).

### 4.6 Block HSIC Lasso is robust to ill-conditioned problems

Single-cell RNA-seq datasets differ from microarray datasets in two ways. First, the number of features is larger, equaling the number of genes in the annotation (> 20 000). Second, the expression matrices are very sparse, due to biological variability (genes actually not expressed in a particular cell) and dropouts (genes whose expression levels have not been measured, usually because they are low, i.e. technical zeroes). In summary, the problem is severely ill-conditioned, and the feature selectors need to deal with this issue. We observed that block HSIC Lasso runs reliably when faced with variations in the data, even on ill-conditioned problems like scRNA-seq. In the different scRNA-seq datasets, LARS was unable to select the requested number of biomarkers in any of the cases, returning always a lower number (Figure S5). mRMR did in all cases. However, the implementation of mRMR that we used crashed while selecting features on the full Villani *et al*. (2017) dataset.

### 4.7 Block HSIC Lasso for biomarker discovery

#### 4.7.1 New biomarkers in mouse hippocampus scRNA-seq

To study the potential of block HSIC lasso for biomarker discovery in scRNA-seq data, we focused on the mouse hippocampus dataset from Habib *et al*. (2016), as a list of 1 669 known biomarkers for the different cell types is also provided by the authors. We requested block HSIC Lasso, mRMR and LARS to select the best 20 genes for classification of 8 cell types (Table S2). The cell types were four different hippocampal anatomical subregions (DG, CA1, CA2 and CA3), glial cells, ependymal, GABAergic and unidentified cells.

The overlap between the genes selected by different algorithms was empty. We compared the selected genes to the known biomarkers. Out of the 20 genes selected by mRMR, 14 are known biomarkers, a number that goes down to 0 in the case of block HSIC Lasso (Figure S5A). Hence, these 20 genes, which are sufficient for accurately separating the cell types, are potential novel biomarkers. However, we have no reason to believe that HSIC Lasso generally has a higher tendency to return novel genes than other approaches; we merely emphasize that it suggests alternative, statistically plausible biological hypotheses that can be worth investigating.

We therefore evaluated whether the novel genes found by block HSIC Lasso participate in biological functions known to be different between the cell classes. To obtain the biological processes responsible for the differences between classes, we mapped the known biomarkers to GO Biological process categories using the GO2MSIG database (Powell, 2014). Then we repeated the process using the genes selected by the different feature selectors, and compared the overlap between them. The overlap between the different techniques increases when we consider the biological process instead of specific genes (Figure S5B). Specifically, one biological process term that is shared between mRMR and block HSIC Lasso, “Adult behaviour” (associated to *Sez6* and *Klhl1*, respectively), is clearly related to hippocampus function. This reinforces the notion that the selected genes are relevant for the studied phenotypes.

Then we focused on potential biomarkers and biologically interesting molecules among those genes selected by block HSIC Lasso. As it is designed specifically to select non-redundant features, often-used GO enrichment analyses are not meaningful: we expect genes belonging to the same GO annotation to be correlated, and HSIC lasso should not accumulate them. Among the top 5 genes, 2 mapped to a biological processes known to be involved: the aforementioned *Klhl1* and *Pou3f1* (related to Schwann cell development). *Klhl1* is a gene expressed in 7 of the studied cell types and which has been related to neuron development in the past (He *et al*., 2006). *Pouf1* is a transcription factor which in the past has been linked to myelination, and neurological damage in its absence (Jaegle *et al*., 1996). The only gene among the top 5 that was expressed exclusively in one of the clusters is the micro RNA *Mir670*, expressed exclusively in CA1. According to miRDB (Wong and Wang, 2015), *Mir670* top predicted target of its 3’ arm is *Pcnt*, which is involved in neocortex development.

#### 4.7.2 GWAS without assumptions on genetic architecture

We applied block HSIC Lasso (*B* = 60, *M* = 1) to three GWA datasets (Section 3.3). It is typical in GWAS to assume a genetic model before performing statistical testing of associations between SNPs and the phenotype. Two common, well-known models are the additive model – the minor allele in homozygosity has twice the effect as the minor allele in heterozygosity – and the dominant model – any number of copies of the minor allele have a phenotypic outcome. Using nonlinear models such as block HSIC Lasso to explore the relationship between SNPs and outcome is attractive since no assumptions are needed on how individual SNPs affect the trait. The only assumption is that the phenotype can be explained by a combination of main effects, as block HSIC Lasso does not account for epistasis. On top of that, by penalizing the selection of redundant features, block HSIC Lasso avoids selecting multiple SNPs in high linkage disequilibrium.

In our experiments, we selected 10 SNPs with block HSIC Lasso for each of the three phenotypes. These are the SNPs that best balance high relatedness to the phenotype and not giving redundant information, be it through linkage disequilibrium or through an underlying shared biological mechanism. We compared these SNPs to those selected by the univariate statistical tests implemented in PLINK 1.9 (Chang *et al*., 2015). Some of them explicitly account for nonlinearity by considering dominant and recessive models of inheritance. The number of SNPs that were positive in at least one test were disparate between the studied phenotypes: all 10 in T1D, 5 in RA, and only 2 in T2D.

Specifically, we compared the genome-wide genotypic p-values to the SNPs selected by block HSIC Lasso (Figure 4). In T1D, block HSIC Lasso selected SNPs among those with the most extreme p-values. However, not being constrained by a conservative p-value threshold, block HSIC Lasso selects five and eight SNPs in RA and T2D, respectively, with non-Bonferroni significant p-values when they improve classification accuracy Interestingly, one of these SNPs can be physically mapped to PFKM (Keildson *et al*., 2014), a gene previously identified in genome-wide studies of T2D. The selected SNPs are scattered all across the genome, displaying the lack of redundancy between them. This strategy gives a more representative set of SNPs than other approaches common in bioinformatics, like selecting the smallest 10 p-values.

**Fig. 4.**
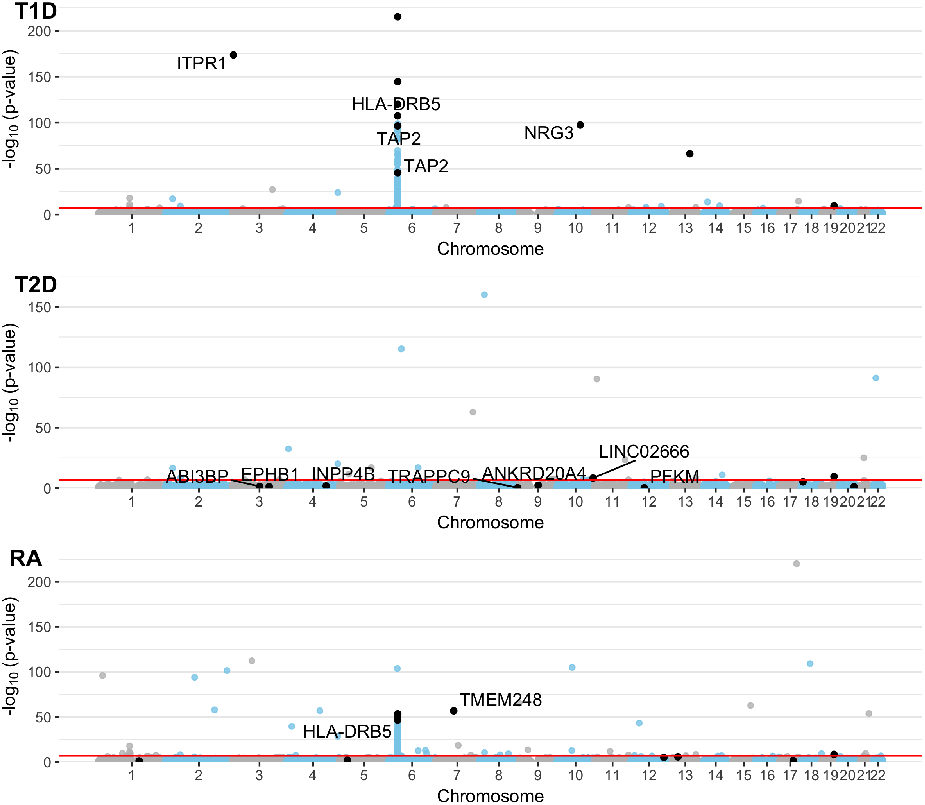
Manhattan plot of the GWA datasets using p-values from the genotypic test. A constant of 10^−220^ was added to all p-values to allow plotting p-values of 0. SNPs in black are the SNPs selected by block HSIC Lasso (*B* = 20), 10 per phenotype. When SNPs are located within the boundaries of a gene (±50 kb), the gene name is indicated. The red line represents the Bonferroni threshold with *α* = 0.05.

## 5 Discussion

In this work, we presented block HSIC Lasso, a nonlinear feature selector. Block HSIC Lasso retains the properties of HSIC Lasso while extending its applicability to larger datasets. Among the attractive properties of block HSIC Lasso we find, first, its ability to handle both linear and nonlinear relationships between the variables and the outcome. Second, block HSIC Lasso has a convex formulation, ensuring that a global solution exists, and that it is accessible. Third, the HSIC score can be accurately estimated, as opposed to other measures of nonlinearity like mutual information. Fourth, block HSIC Lasso’s memory consumption scales linearly with respect to both the number of features and the number of samples. In addition, block HSIC Lasso can be easily adapted to different problems via different kernel functions that better capture similarities in new datasets. Lastly, block HSIC Lasso can be adjusted for covariates known to affect the outcome, which helps removing confounding effects from the analysis. Due to all these properties, we show how block HSIC Lasso outperforms all other algorithms in the tested conditions.

Block HSIC Lasso can be applied to different kinds of datasets. As other nonlinear methods, block HSIC Lasso is particularly useful when we do not want to make strong assumptions about how the causal variables relate to the outcome. Thanks to the advantages mentioned above, HSIC Lasso and block HSIC Lasso tend to outperform other state-of-the-art approaches in terms of both causal features retrieval in simulated data, and classification accuracy on real-world datasets.

Whereas the Lasso is limited to selecting at most as many features as there are available samples (*n*), for block HSIC Lasso the limitation is *nBM*. Hence, even if the number of samples is small, block HSIC Lasso can be used to select a larger number of features. If *nBM* is still limiting, one could replace the *ℓ*_1_ regularization with an elastic-net regularization. However, in most cases, we expect block HSIC Lasso to be used to select a small number of features.

Regarding its potential in bioinformatics, we applied block HSIC Lasso to images, microarrays, single-cell RNA-seq and GWAS. The two latter involve thousands of samples, making it unfeasible to run vanilla HSIC Lasso on a regular server because of its memory requirements. The selected biomarkers are biologically plausible, agree with the outcome of other methods, and provide a good classification accuracy when used to train a classifier. Such a ranking is useful, for instance, when selecting SNPs or genes to assay in *in vitro* experiments.

Block HSIC Lasso’s main drawback is the memory complexity, markedly lower than in vanilla HSIC Lasso but still *O*(*dnB*). Memory issues might appear in low-memory servers in cases with a large number of samples *n*, of features *d*, or both. However, through our work on GWA datasets, the largest type of dataset in bioinformatics, we show that working on these datasets is feasible. Another drawback, which block HSIC Lasso shares with the other nonlinear methods, is their black box nature. Block HSIC Lasso looks for biomarkers which, after an unknown, nonlinear transformation, would allow a linear separation between the samples. Unfortunately, we cannot access this transformed space and explore it, which makes the results hard to interpret.

## References

Burton, P. R. et al. (2007). Genome-wide association study of 14,000 cases of seven common diseases and 3,000 shared controls. Nature, 447(7145), 661–678.

Chang, C. C. et al. (2015). Second-generation PLINK: rising to the challenge of larger and richer datasets. GigaScience, 4, 7.

Clarke, R. et al. (2008). The properties of high-dimensional data spaces: implications for exploring gene and protein expression data. Nature Reviews Cancer, 8(1), 37–49.

Cover, T. M. and Thomas, J. A. (2006). Elements of Information Theory. John Wiley & Sons, Inc., Hoboken, NJ, USA, 2nd edition.

Ding, C. and Peng, H. (2005). Minimum redundancy feature selection from microarray gene expression data. Journal of Bioinformatics and Computational Biology, 03(02), 185–205.

Efron, B. et al. (2004). Least angle regression. The Annals of statistics, 32(2), 407–499.

Fujishige, S. (2005). Submodular functions and optimization, volume 58. Elsevier.

Gretton, A. et al. (2005). Measuring statistical dependence with Hilbert-Schmidt norms. In International conference on algorithmic learning theory (ALT), pages 63–77.

Haber, A. L. et al. (2017). A single-cell survey of the small intestinal epithelium. Nature, 551(7680), 333–339.

Habib, N. et al. (2016). Div-Seq: Single-nucleus RNA-Seq reveals dynamics of rare adult newborn neurons. Science (New York, N.Y.), 353(6302), 925–8.

He, Y. et al. (2006). Targeted deletion of a single Sca8 ataxia locus allele in mice causes abnormal gait, progressive loss of motor coordination, and Purkinje cell dendritic deficits. The Journal of neuroscience: the official journal of the Society for Neuroscience, 26(39), 9975–82.

Jaegle, M. et al. (1996). The POU factor Oct-6 and Schwann cell differentiation. Science (New York, N.Y.), 273(5274), 507–10.

Johnstone, I. M. and Titterington, D. M. (2009). Statistical challenges of high-dimensional data. Philosophical transactions. Series A, Mathematical, physical, and engineering sciences, 367(1906), 4237–53.

Keildson, S. et al. (2014). Expression of phosphofructokinase in skeletal muscle is influenced by genetic variation and associated with insulin sensitivity. Diabetes, 63(3), 1154–65.

Li, J. et al. (2018). Feature selection: A data perspective. ACM Computing Surveys (CSUR), 50(6), 94.

Mairal, J. et al. (2010). Online learning for matrix factorization and sparse coding. Journal of Machine Learning Research, 11, 19–60.

Peng, H. (2005). mrmr. http://home.penglab.com/proj/mRMR/.

Peng, H. et al. (2005). Feature selection based on mutual information: Criteria of max-dependency, max-relevance, and min-redundancy. IEEE Transactions on Pattern Analysis and Machine Intelligence, 27, 1226–1237.

Powell, J. A. C. (2014). GO2MSIG, an automated GO based multispecies gene set generator for gene set enrichment analysis. BMC bioinformatics, 15, 146.

Ravikumar, P. et al. (2009). Sparse additive models. Journal of the Royal Statistical Society: Series B (Statistical Methodology), 71(5), 1009–1030.

Schölkopf, B. and Smola, A. J. (2002). Learning with Kernels. MIT Press, Cambridge, MA.

Song, L. et al. (2012). Feature selection via dependence maximization. Journal of Machine Learning Research, 13, 1393–1434.

Tibshirani, R. (1996). Regression shrinkage and selection via the Lasso. Journal of the Royal Statistical Society, Series B, 58(1), 267–288.

van Dijk, D. et al. (2018). Recovering Gene Interactions from Single-Cell Data Using Data Diffusion.

Villani, A.-C. et al. (2017). Single-cell RNA-seq reveals new types of human blood dendritic cells, monocytes, and progenitors. Science (New York, N.Y.), 356(6335), 925–8.

Walters-Williams, J. and Li, Y. (2009). Estimation of mutual information: A survey. In P. Wen, Y. Li, L. Polkowski, Y. Yao, S. Tsumoto, and G. Wang, editors, Rough Sets and Knowledge Technology, pages 389–396, Berlin, Heidelberg. Springer Berlin Heidelberg.

Wong, N. and Wang, X. (2015). miRDB: an online resource for microRNA target prediction and functional annotations. Nucleic acids research, 43(Database issue), D146–52.

Yamada, M. et al. (2014). High-dimensional feature selection by feature-wise kernelized lasso. Neural computation, 26(1), 185–207.

Yamada, M. et al. (2018). Ultra high-dimensional nonlinear feature selection for big biological data. IEEE Transactions on Knowledge and Data Engineering, 30(7), 1352–1365.

Zhang, Q. et al. (2018). Large-scale kernel methods for independence testing. Statistics and Computing, 28(1), 113–130.

