## Supplementary File 1 for "Block HSIC Lasso: model-free biomarker detection for ultra-high dimensional data"

### Supplementary file 1: Advantages of block HSIC Lasso over existing nonlinear feature selection methods

March 22, 2019

Compared to linear feature selection algorithms, there are a relatively small number of nonlinear feature selection methods for high-dimensional data. The widely used Sparse Additive Models (SpAM) [Ravikumar *et al.*, 2009] can be useful to find a set of features if the output is the result of an additive function. However, in biology this assumption might be too strong, so SpAM may not be able to find the relevant features. An alternative approach are association-based approaches [Balasubramanian *et al.*, 2013, Fan and Lv, 2008]. They first reduce the number of features by using a statistic (e.g. mutual information) and then select the final set of important features from the screened feature sets. This two-step approach is useful in practice. However, since screening-based approaches use a measure of similarity between features and outcome to screen features, it can select redundant features. Recently, several nonlinear feature selection algorithms were proposed including the gradient boosted feature selection [Xu *et al.*, 2014] and kernel based methods [Gregorová *et al.*, 2018, Tan *et al.*, 2014].

The key advantage of the proposed algorithm, block HSIC Lasso, is that it is a versatile nonlinear feature selection algorithm. More specifically, by using different kernels it can not only handle continuous and discrete inputs, but also multi-variate and multi-class outcomes. Moreover, it can select features from ultra high-dimensional and large-scale data in a few hours of computation.

#### References


Balasubramanian, K. *et al.* (2013). Ultrahigh dimensional feature screening via RKHS embeddings. In *Artificial Intelligence and Statistics (AISTATS)*, pages 126–134.

Fan, J. and Lv, J. (2008). Sure independence screening for ultrahigh dimensional feature space. *Journal of the Royal Statistical Society: Series B (Statistical Methodology)*, **70**(5), 849–911.

Gregorová, M. *et al.* (2018). Large-scale nonlinear variable selection via kernel random features. In *Joint European Conference on Machine Learning and Knowledge Discovery in Databases (ECMLPKDD)*, pages 177–192.

Ravikumar, P. *et al.* (2009). Sparse additive models. *Journal of the Royal Statistical Society: Series B (Statistical Methodology)*, **71**(5), 1009–1030.

Tan, M. *et al.* (2014). Towards ultrahigh dimensional feature selection for big data. *Journal of Machine Learning Research*, **15**, 1371–1429.

Xu, Z. *et al.* (2014). Gradient boosted feature selection. In *KDD*.
