## Supplementary Figures for "Block HSIC Lasso: model-free biomarker detection for ultra-high dimensional data"

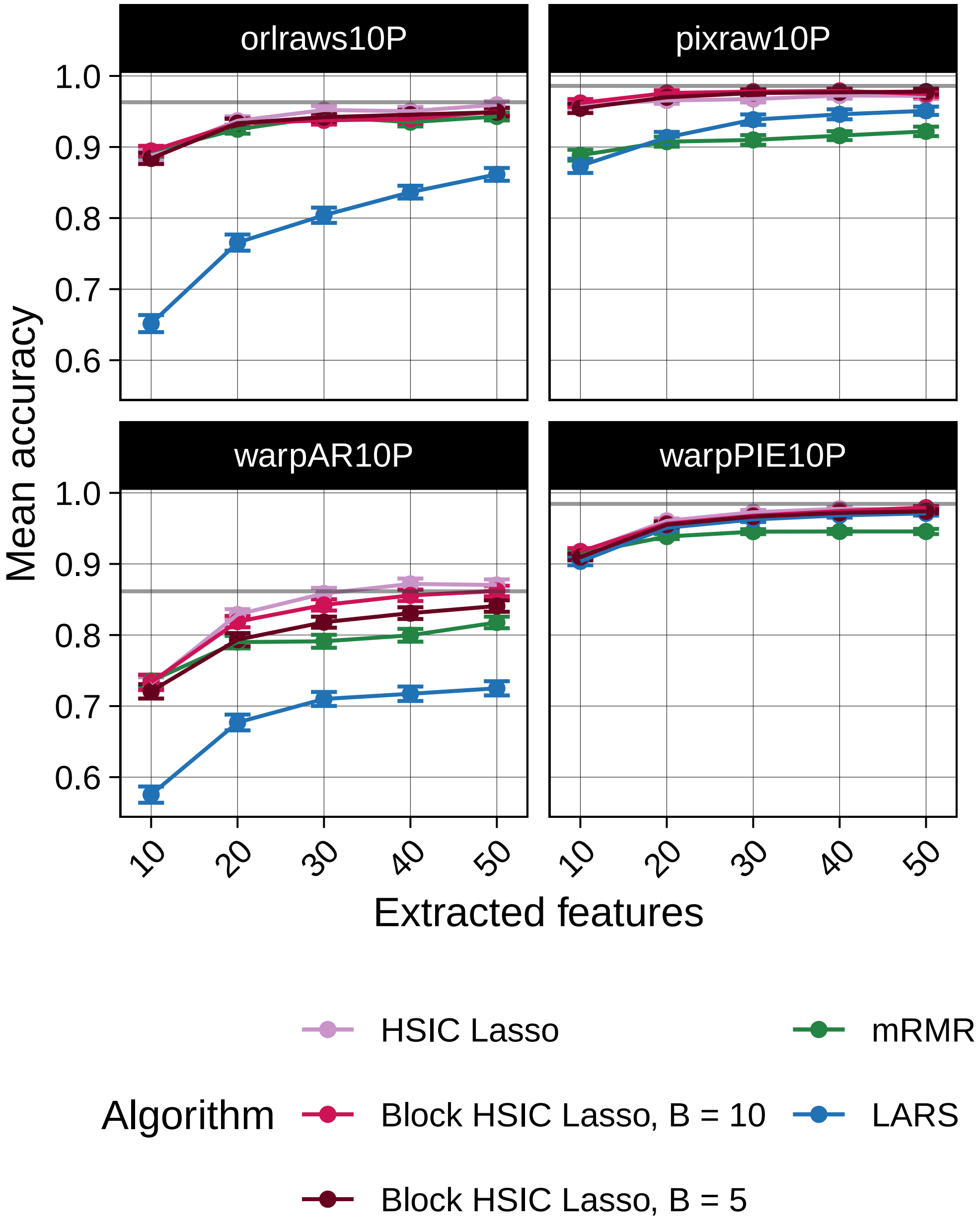


Figure S1: Prediction accuracy of a random forest classifier trained on features selected by different algorithms in four image datasets. The gray line represents the mean accuracy of 10 classifiers trained on all the dataset.

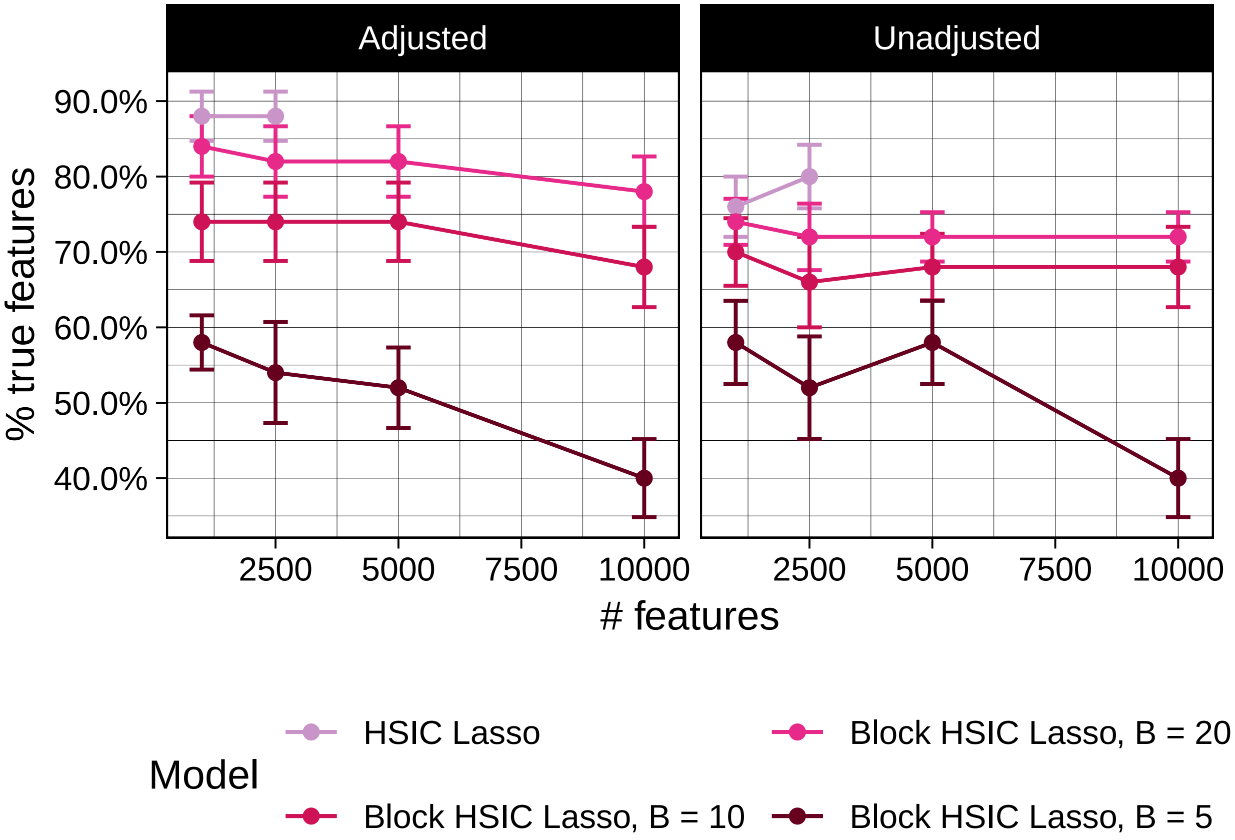


Figure S2: Correct number of causal features detected by (block) HSIC Lasso, with, and without adjustment for covariates.

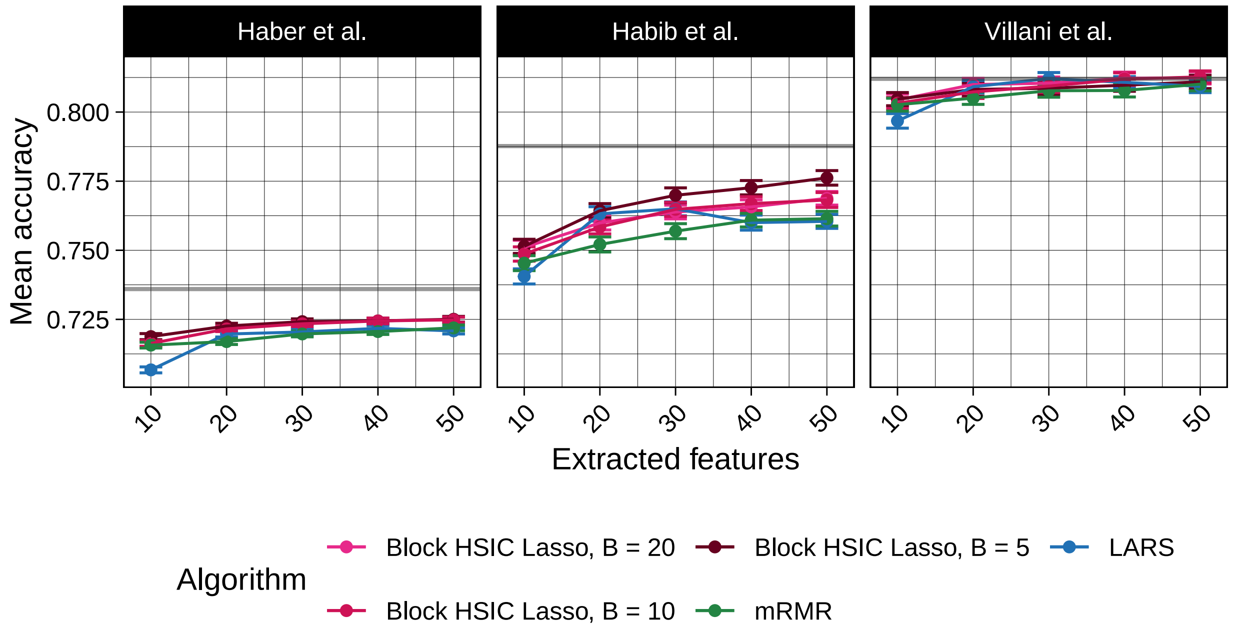


Figure S3: Classification accuracy of a random forest classifier on three single-cell RNA-seq experiments (Section 3.3) after feature selection. The gray line represents the mean accuracy of 10 classifiers trained on all the dataset.

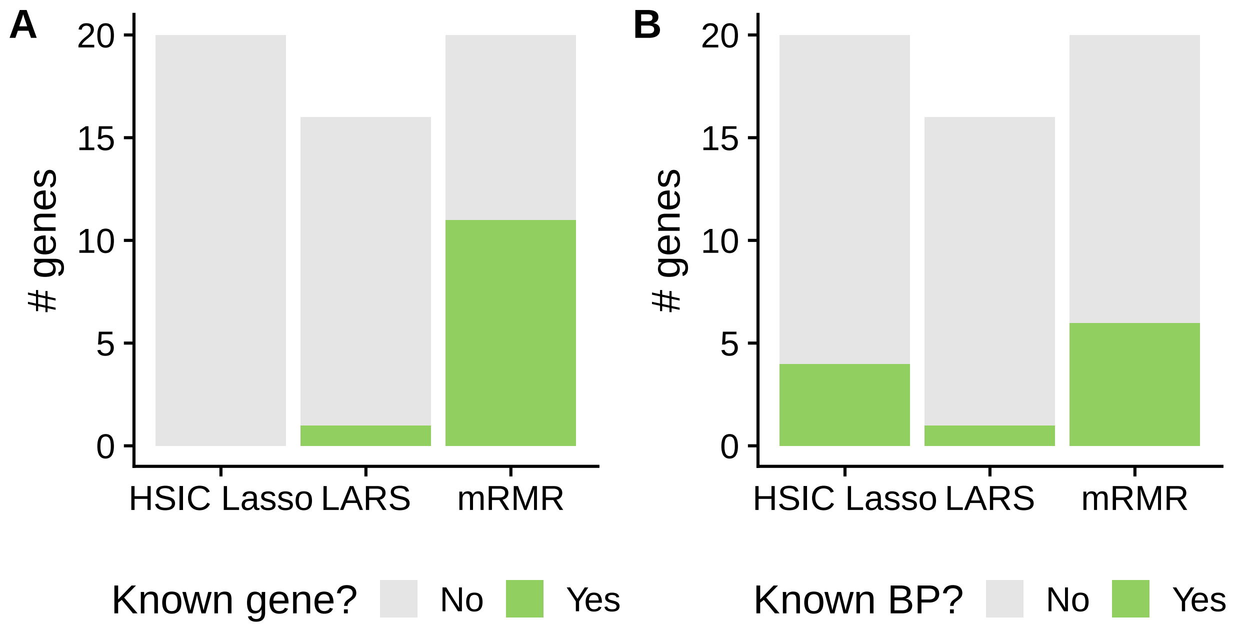


Figure S4: Genes selected by different feature selection algorithms on single-cell RNA-seq data from mouse hippocampus (Habib et al., 2016). A. Count of genes that are (not) known biomarkers. B. Count of genes that share (not) a GO Biological process with a known biomarker.

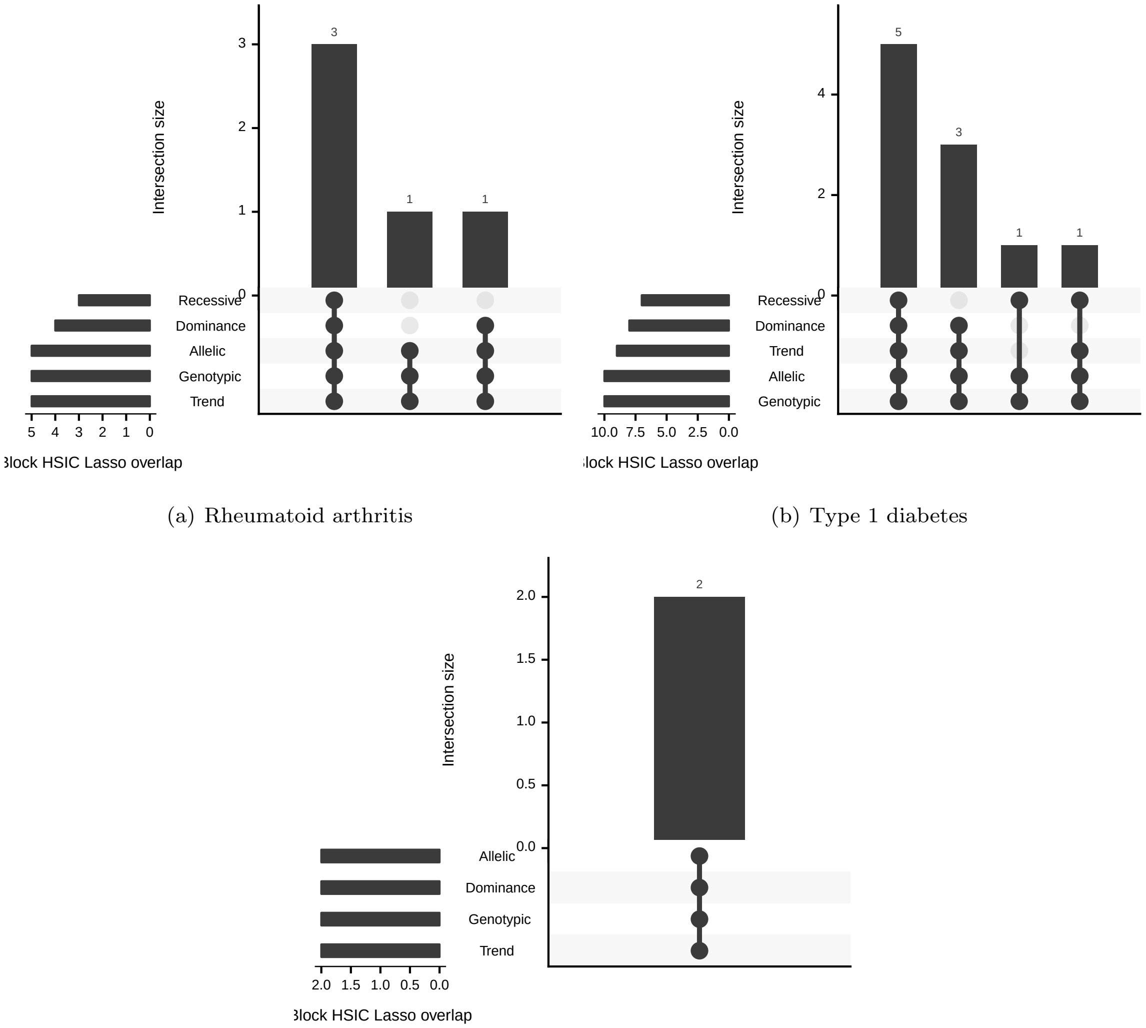


(a) Rheumatoid arthritis

(b) Type 1 diabetes

(c) Type 2 diabetes

Figure S5: Subset of block HSIC Lasso-detected biomarkers that were detected ( $p < p_{\text{bonferroni}}$ ) in several univariate statistical tests.

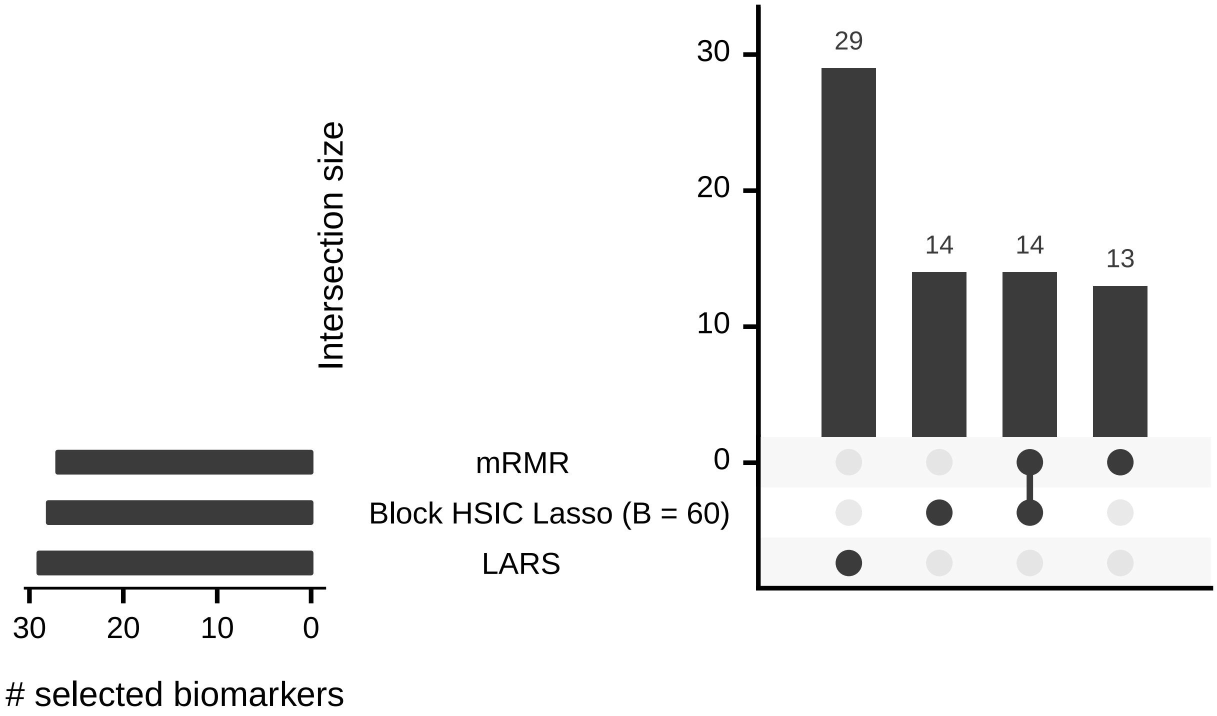


Figure S6: Overlap between the pooled SNPs selected by three feature selectors (block HSIC Lasso, mRMR and LARS) on three GWA datasets. We asked each method to select 10 features in each dataset. The horizontal histogram represents the total number of SNPs selected by each algorithm. The vertical histogram represents the size of the overlap between SNP sets (14 SNPs are selected by both block HSIC Lasso and mRMR; 14 were exclusively selected by block HSIC Lasso or mRMR and so on).
